## Supplementary Information for "Optimized CRISPR tools and site-directed transgenesis in *Culex quinquefasciatus* mosquitoes for gene drive development"

##### Supplementary Figure 1

**a.**

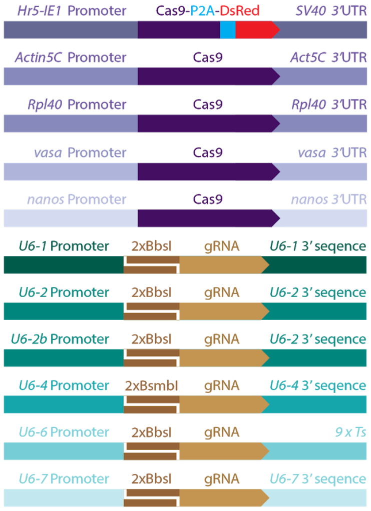


| Promoter | Strain | Plasmid | Accession |
|---|---|---|---|
| Baculovirus IE1 | n/a | pVMG225_IE1>Cas9-P2A-DsRed | To be deposited |
| CPIJ009808 | CA | pVMG173_*Cq-Actin5c*>Cas9 | To be deposited |
| CPIJ002413 | CA | pVMG193_*Cq-Rpl40*>Cas9 | To be deposited |
| CPIJ009286 | CA | pVMG213_*Cq-vasa*>Cas9 | To be deposited |
| CPIJ011551 | CA | pVMG212_*Cq-nanos*>Cas9 | To be deposited |
| CPIJ039653 | AL | pVMG146_*Cq-U6:1*>2xBbsI-gRNA | To be deposited |
| CPIJ039728 | AL | pVMG147_*Cq-U6:2*>2xBbsI-gRNA | To be deposited |
| CPIJ039728 | CA | pVMG217_*Cq-U6:2b*>2xBbsI-gRNA | To be deposited |
| CPIJ039801 | CA | pVMG218_*Cq-U6:4*>2xBsmbI-gRNA | To be deposited |
| CPIJ039596 | AL | pVMG149_*Cq-U6:6*>2xBbsI-gRNA | To be deposited |
| CPIJ040693 | CA | pVMG164_*Cq-U6:7*>2xBbsI-gRNA | To be deposited |

**b.**

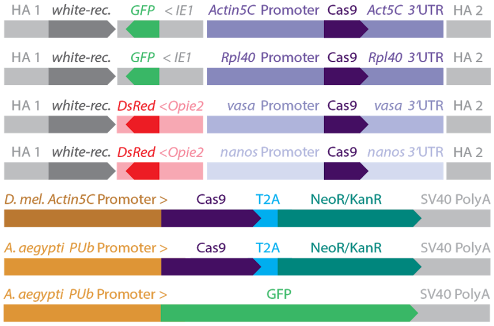


| Plasmid | Accession |
|---|---|
| pVMG177_*CqActin5c*>Cas9_wHAs | To be deposited |
| pVMG220_*CqRpl40*>Cas9_wHAs | To be deposited |
| pVMG236_*CqVasa*>Cas9_wHAs | To be deposited |
| pVMG235_*CqNanos*>Cas9_wHAs | To be deposited |
| p*Dm-Act5C*>Cas9-T2A-Neo | To be deposited |
| p*Aae*-PUb>Cas9-T2A-Neo | To be deposited |
| pSL1180-HDR-*Aae*-PUb>GFP-NLS | To be deposited |

**c.**

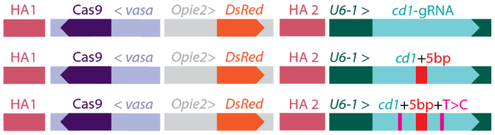


| Plasmid | Accession |
|---|---|
| pVMG252_*Cq-vasa*>Cas9_cdHAs_O | To be deposited |
| pVMG280_*Cq-vasa*>Cas9_cdHAs_L | To be deposited |
| pVMG277_*Cq-vasa*>Cas9_cdHAs_L+M | To be deposited |

**d.**

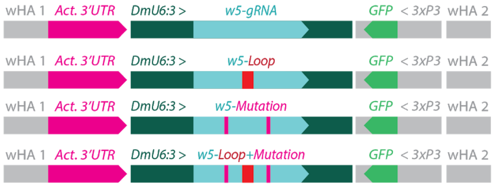


| Plasmid | Accession |
|---|---|
| pVMG109_CC-*U6:3-w5*_GFP_wHAs_O | To be deposited |
| pVMG221_CC-*U6:3-w5*_GFP_wHAs_L | To be deposited |
| pVMG222_CC-*U6:3-w5*_GFP_wHAs_M | To be deposited |
| pVMG274_CC-*U6:3-w5*_GFP_wHAs_L+M | To be deposited |

**Supplementary Figure 1 - Reagents that have been built for the experiments in this manuscript. (a)** Transgenes built for the expression of Cas9 and gRNA. The gRNAs constructs have been further modified by adding the gRNA target sequences or the scaffold variants, such further edits are not reported in this table. Next to the construct structure are reported the genes from which genomic sequences were amplified and the laboratory line from which genomic DNA and PCR amplification was run. **(b)** Variation of the Cas9 constructs used to test activity in cells and in embryos. These constructs were tested instead of the simpler ones as we originally intended to target the white gene for transgenesis. Additionally we tested two exogenous promoters, *Actin5C* from *Drosophila melanogaster* (*D. mel.*) and *PUb* from *Aedes aegypti*. (**c**) HR template plasmids built for transgenesis of the vasa-Cas9 cassette in *Culex quinquefasciatus.* (**d**) Constructs used to evaluate transgenesis and gene drive performance in *Drosophila melanogaster.* **Accession numbers will be added ahead of publication.**

**Supplementary Figure 2**

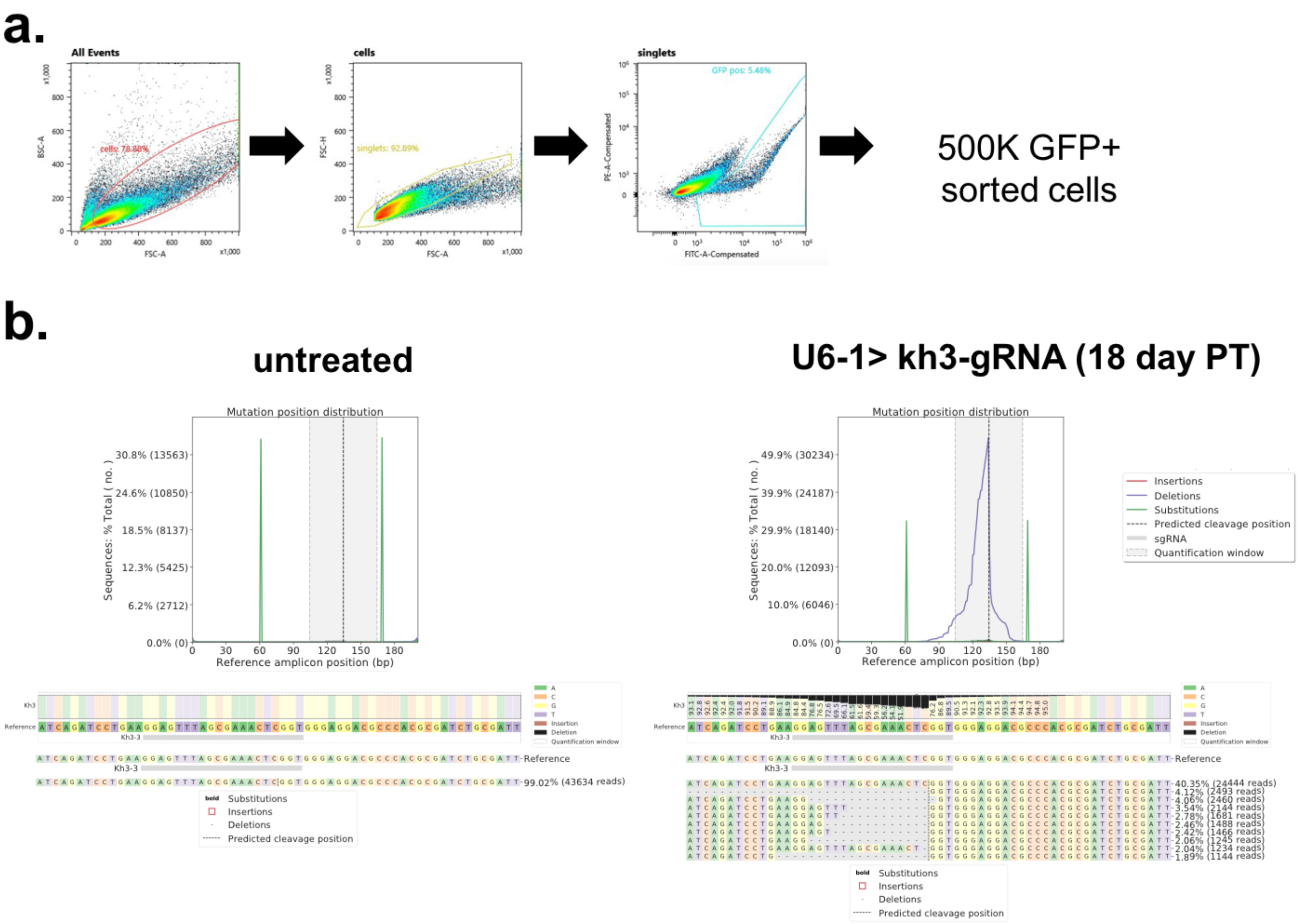


**Supplementary Figure 2 - FACS gating strategy and CRISPResso2 analysis output. (a)** Gating strategy used for FACS sorting of GFP expressing cells. **(b)** Example of CRISPResso2 analysis output for untreated and kh3-gRNA targeted sample. The graph shows all types of mutation occurring in the kh3 target amplicon represented by category; The quantification window is highlighted in gray and the broken line represents the exact cutting site (-3bp from PAM). Below is represented the % of nucleotide substitutions or deletions around the cutting site. At the bottom, a 60bp sequence window centered on the cutting site shows mutated allele sequences and their relative % abundance.

**Supplementary Figure 3**

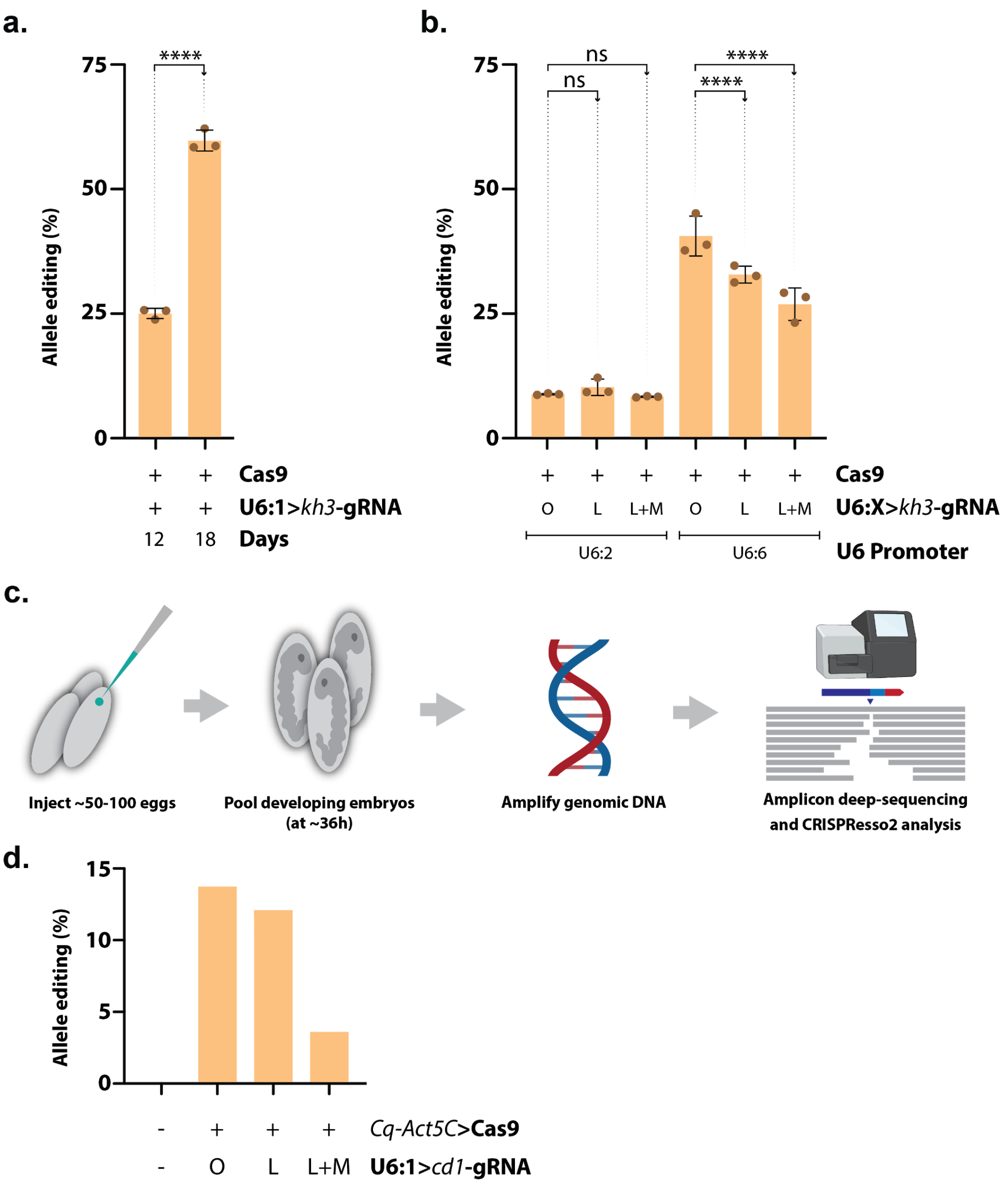


**Supplementary Figure 3 - Additional testing of the *Culex quinquefasciatus* reagents in cells and *in vivo*.** **(a)** Histogram representing editing efficiency (%) of *kh3* target locus in Hsu cells when co-transfected with a mixture of *Cq-Actin5C*>Cas9 paired to U6:1>*kh3*-gRNA and analyzed 12 days (left bar) or 18 days (right bar) post-transfection. (**b**) Editing efficiency (%) at *kh3* target locus in Hsu cells when transfected with a mixture of *Cq-Actin5C*>Cas9 paired either to U6:2 or U6:6 *pol-III* promoters expressing the *kh3*-gRNA with different gRNA scaffold variants. Histogram bars represent the mean, error bars and dots represent SD and distribution of 3 biological replicates. Statistical comparisons were generated with a two way ANOVA multiple comparison of the mean, with Tukey's corrections. **** = $P_{tukey} < 0.0001$, ns = $P_{tukey} > 0.05$. (**c**) Modified version of our protocol in **Fig. 1d** that we used to evaluate *in vivo* the allele editing rates of the gRNA variants. The *Cq-Actin5C*>Cas9 plasmid was co-injected with U6:1>*cd1*-gRNA with gRNA scaffold variants, then we collected DNA from all developing embryos at ~36h post injection for the subsequent deep sequencing analysis. (**d**) Edited allele percentages observed at the *cd1* site, for the scaffold variants evaluated. The control was generated from a pool of uninjected eggs. (**b, d**) O = “Original”, L = “Loop”, L+M = “Loop+Mutation”.

**Supplementary Figure 4**

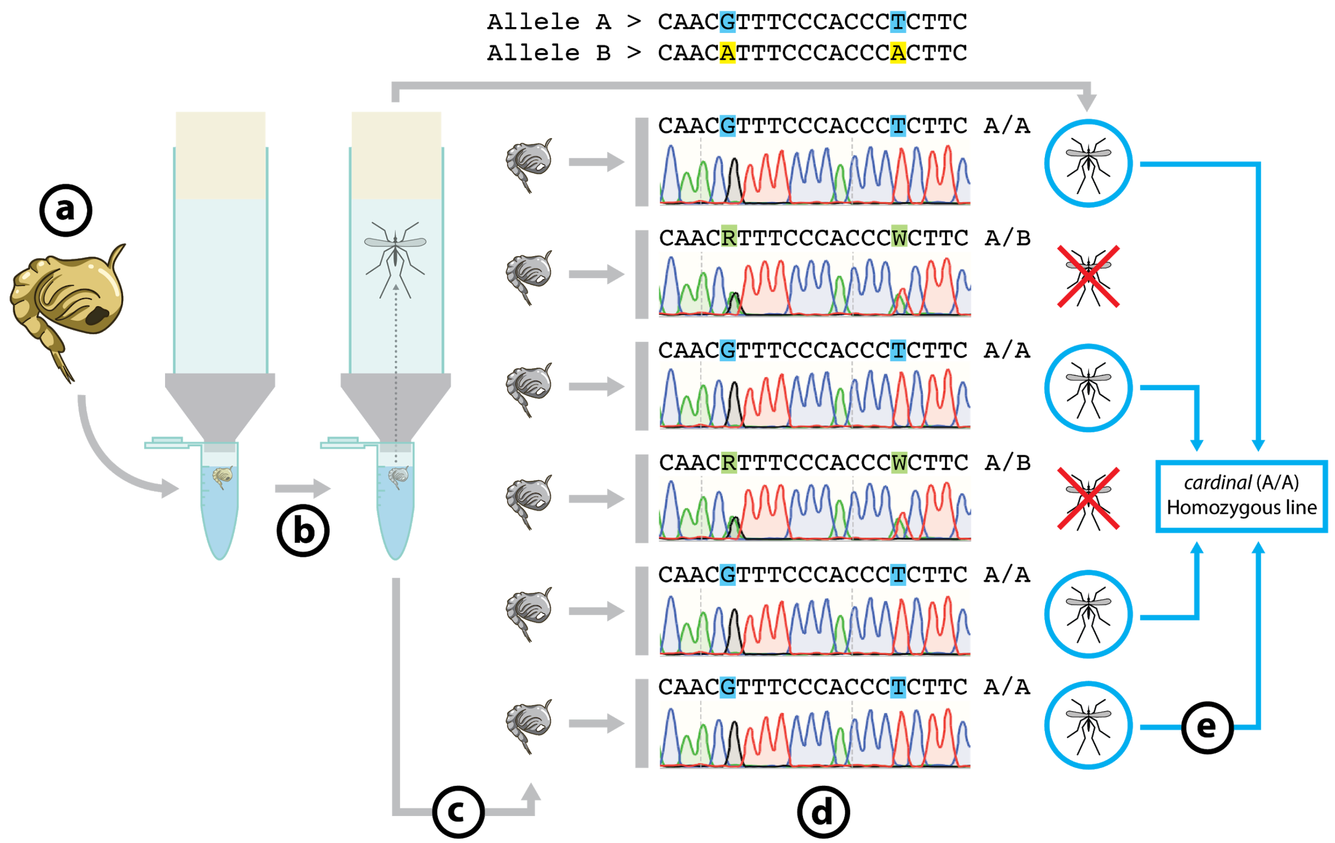


**Supplementary Figure 4 - Generation of a *Culex quinquefasciatus* line isogenic at the *cardinal* locus. (a)** Pupae about to hatch were collected and individually deposited in different Eppendorf tubes with about 1mL of water. Each tube was then coupled with a capped plastic *Drosophila* vial with the bottom sawed off, using a 3D-printed adaptor. **(b)** After ~24 hours the hatched mosquitoes were maintained in the respective tubes, while **(c)** the pupal case was removed and used for PCR amplification of the *cardinal* locus. **(d)** Resulting amplicons were used for Sanger sequence analysis of the *cardinal* locus and evaluate the individuals' genotype (A/A, A/B or B/B; we did not recover any B/B animals suggesting lethality associated with this allele). **(e)** All A/A male and female mosquitoes were pooled in a cage to generate a line homozygous for the cardinal-A allele, about ~24h after hatching. Note that the mosquitoes were able to survive overnight within the collection vials without needing a source of food or water as they were kept in a high-humidity environment.

#### Supplementary Figure 5

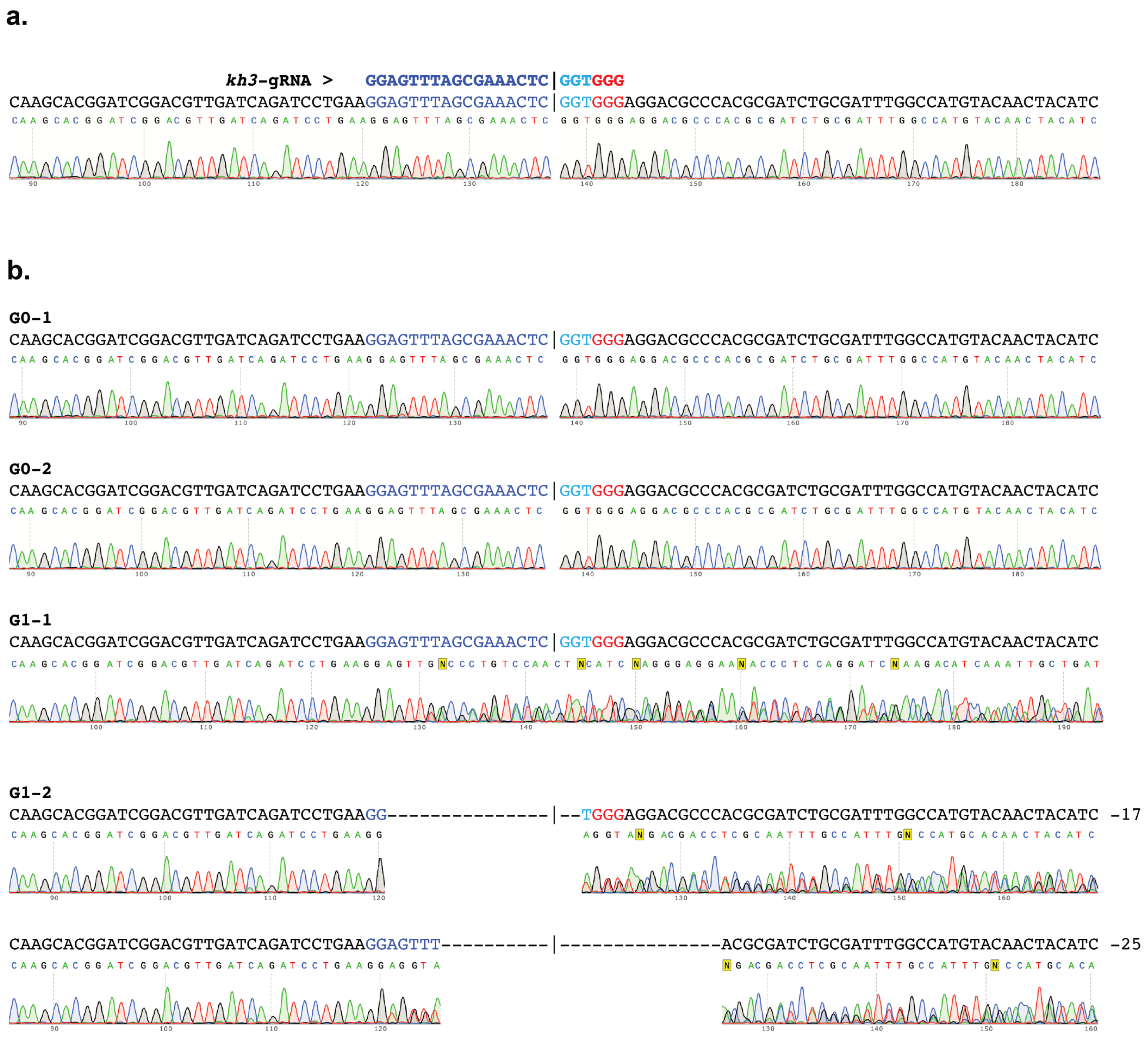


**Supplementary Figure 5 - Sequencing of G0 and G1 animals from the Cas9-line validation experiment. (a)** Wild-type locus, showing the location of the gRNA target in blue and the PAM in red. A vertical bar indicates the location of the cut site. **(b)** Sanger sequences traces obtained from two G0 injected mosquitoes showing no apparent cutting; and two traces obtained from two G1 animals displaying a *kh-* (white eye) phenotype. For the bottom animal (G1-2) we were able to disentangle the two indels present by using the ICE analysis software by Synthego (Synthego Performance Analysis, ICE Analysis. 2019. v2.0. Synthego; 15 Oct. 2020), with a Sanger sequencing read performed on wild-type as a control; we have aligned the same sequence trace at the positions corresponding with each indel.

**Supplementary Figure 6**

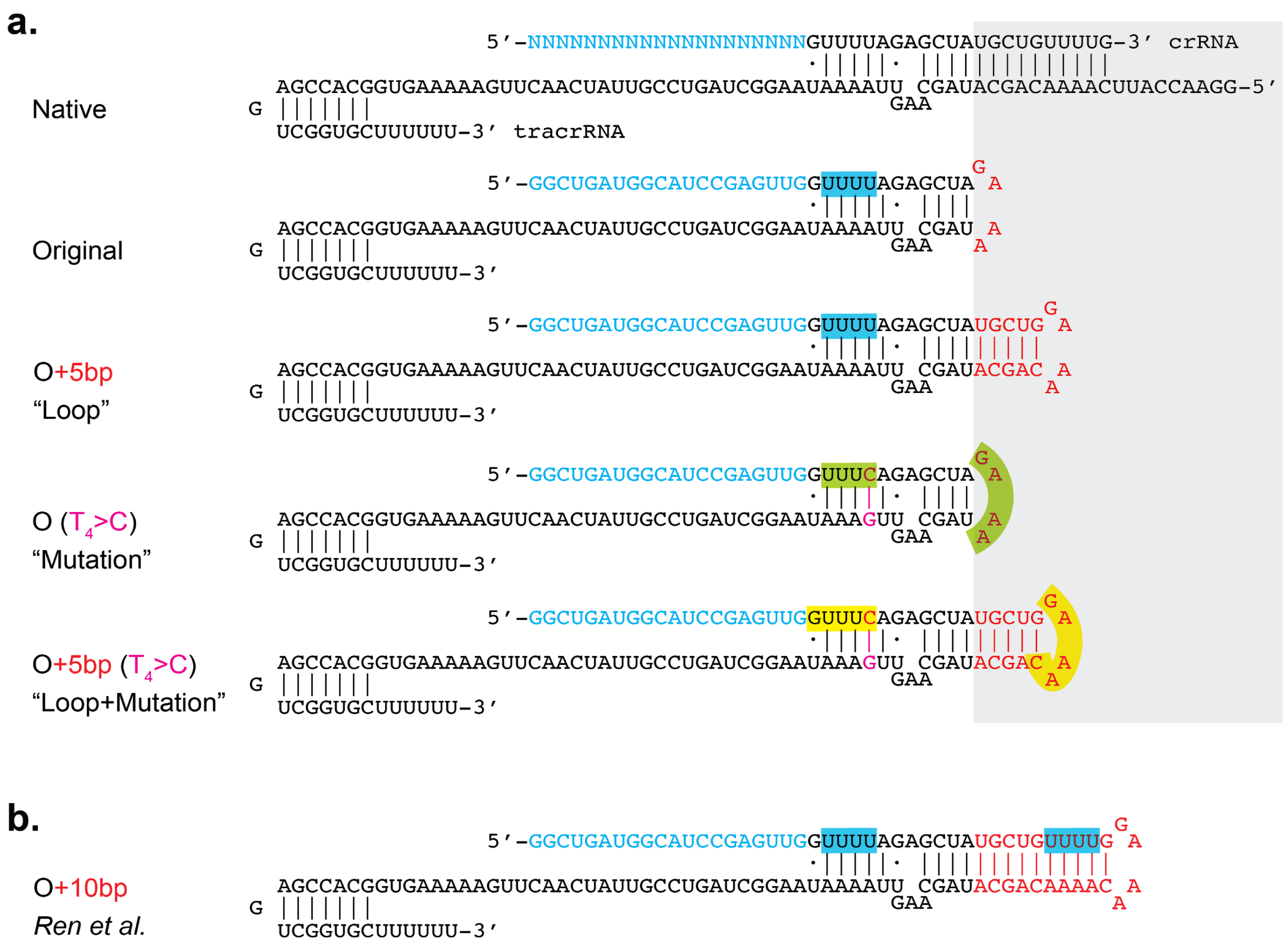


**Supplementary Figure 6 - Potential mispairing of the scaffold variants containing the $T_4$>C mutation.** (**a**) Shows the different gRNA variants tested in this study and their RNA base-pairing compared to the native crRNA/tracrRNA couple. For the gRNAs containing the $T_4$>C mutation, we identified a potential pairing between the sequence present in the first loop "GAAA" with a sequence present at the beginning of the gRNA scaffold UUUC, highlighted in green. When the mutation is combined with the loop, the added sequence introduces an even longer potential base-pairing of the GAAAC with the GUUUC sequence, highlighted in yellow. (**b**) Modified scaffold tested in *Ren et al.* with an additional 10bp in the loop. Highlighted in blue are the sequences, present as Ts in the transgene that could potentially trigger premature stop of transcription.

#### Supplementary Tables

##### Supplementary Table 1 - Additional injection conditions with supplemented Cas9 protein and/or plasmid mixture.

| Plasmid | Round of Injection | Cas9 Source | | Injected G0 | | | Efficiency (per G0 germline)* | | Overall efficiency (out of total G1s)# | |
|---|---|---|---|---|---|---|---|---|---|---|
| | | Protein | Plasmid Mix | Injected eggs | Adult survivors | Survival (%) | Cutting *cd-/cd-* (%) | Transgenic DsRed+ (%) | Cutting *cd-/cd-* (%) | Transgenic DsRed+ (%) |
| ***cd1*-original** | 1 | + | + | 320 | 36 | 11.25% | 0/36 (0.00%) | 0/36 (0.00%) | 0/398 (0.00%) | 0/398 (0.00%) |
| ***cd1*-loop** | 1 | +^ | + | 460 | 9 | 1.96% | **3**/9 (33.33%) | **2**/9 (22.22%) | **48**/431 (11.14%) | **39**/431 (9.05%) |
| | 2 | + | + | 412 | 16 | 3.88% | **1**/16 (6.25%) | 0/16 (0.00%) | **15**/804 (0.19%) | 0/804 (0.00%) |
| | 1 | + | - | 280 | 13 | 4.64% | 0/13 (0.00%) | 0/13 (0.00%) | 0/982 (0.00%) | 0/982 (0.00%) |
| | 1 | - | + | 550 | 55 | 10.00% | 0/55 (0.00%) | 0/55 (0.00%) | 0/2666 (0.00%) | 0/2666 (0.00%) |

* The G0 germline cutting and transgenesis efficiencies were calculated as numbers of independent pools that produce either *cd-/cd-* mutant (cutting) or DsRed+ (transgenesis) animals, divided by the total number of crossed G0s. While each pool contains several G0 individuals, in our calculations we assume only one editing event happening in each positive pool, therefore our calculations might underestimate the cutting and transgenesis rates.
### The overall cutting and transgenesis efficiencies were calculated as the number of G1 individuals with either *cd-/cd-* (cutting) or DsRed+ (transgenesis) phenotypes divided by the total number of G1s.
^ In this injection we used an older batch of Cas9 protein which later showed low activity in subsequent studies.

**SUPPLEMENTARY DATA**

**Supplementary Data 1 - Genome editing quantification data.**

Summary of the CRISPResso2 batch analysis on allele editing frequencies performed on deep sequencing data deriving from *in vivo* and *in vitro* experiments. For each sample listed in tab 1 and tab 2 are specified: experimental details including names and ratio (%) of plasmid DNA transfected/injected and the experimental group; The raw CRISPResso allele frequency analysis output; The data extrapolated from the batch analysis to calculate allele editing and plotted in Fig1 and Supplementary Figure 2. The data as described is subdivided in the following file tabs:

1. **Cell line editing quantification**
2. **Embryo editing quantification**
3. **CRISPResso2 parameters:** Includes the running parameters used for all analysis on CRISPResso2 with one example representative of each batch experiment run.

**Supplementary Data 2 - *Culex quinquefasciatus* transgenesis counting data.**

Different injection conditions with G0 survival efficiencies were listed. Raw counting data of the G1 progeny phenotype indicating the cutting/ transgenesis in different conditions. *cd-/cd-* phenotype indicates the cutting and DsRed+ indicated the integration. All G0s were divided as male and female pools and crossed with *cd-/cd-*. Egg rafts from each male pool were scored together, while rafts of each female pool were divided and hatched singly in different batches, which gives more precise evaluation of germline rates. The data is subdivided in the following file tabs:

1. **Figure 2 - Table 1 Data:** The cutting/transgenesis efficiencies in conditions of injecting different HDR template variants (HR+Original gRNA; HR+“Loop”; HR+“Loop+Mutation”) were recorded. The germline and overall efficiencies were calculated.
2. **Supplementary Table 1 Data:** The cutting/transgenesis efficiencies in other conditions by injecting HDR templates supplemented with Cas9 resources (Cas9 protein; Cas9 plasmid mixture; and both) were recorded. The germline and overall efficiencies were calculated.

**Supplementary Data 3 - Cas9 line validation counting data.**

Two experimental conditions of injecting *in-vitro*-transcribed *Kh3*-gRNA or *Kh3*-gRNA plasmid were listed. The embryos of heterozygous Cas9 line were used for injection. The injection conditions and G0 situations were recorded. The Cas9 positive G0s were crossed with *kh-/kh-* mutants to evaluate gRNA activity in G1. The cutting efficiencies were calculated based on numbers of individuals giving *kh-* mutant phenotype in G1s. The data is subdivided in the following file tabs:

1. **Figure 3e Data:** The editing rate observed for either IVT-gRNA or plasmid is calculated.

**Supplementary Data 4 - Effect of the gRNA scaffold with the loop modification on transgenesis efficiency.**

Raw counting data of the G1 progeny phenotypic scoring indicating total females and males screened. All G0 crosses were performed in single-pairs between one injected individual and one wildtype (non-injected) animal. Number of recovered GFP positive (GFP+) and non-GFP (GFP-) individuals from the G1 are indicated. This allows us to precisely evaluate single-germline transgenesis rates in our experimental conditions. The data is subdivided in the following file tabs:

1. **Fig. 5 (w5 original)**: Trangenesis rates after injecting the w5-gRNA element carrying the “Original” gRNA scaffold version. Independent germlines that gave rise to transformants (GFP+) and total germlines analyzed as well as transformation rates are indicated.
2. **Fig. 5 (w5 loop)**: Trangenesis rates after injecting the w5-gRNA element carrying the “loop” gRNA scaffold modification. Independent germlines that gave rise to transformants (GFP+) and total germlines analyzed as well as transformation rates are indicated.
3. **Statistical Analysis.** Test for the Difference in Proportions between w5 “Original” injection vs. w5 “Loop” injection.

**Supplementary Data 5 - Gene drive experiments with different gRNA scaffold variants in *Drosophila melanogaster***.

Raw counting data of the F2 progeny phenotypic scoring indicating females and males recovered. Red marker (DsRed+), green marker (GFP+), both fluorophores (both), no fluorescence (none), wild-type eye (w+), white-eye (w-), and mosaic eyes (mosaic) were scored in order to track Cas9 (red marker) and the gRNA (green marker) transgenes, as well as other outcomes of the cross. Transgene inheritance rates in the F2 progeny for each specific tube (marked as “F1 Cross” in the table) were calculated by combining data from males and females. Average inheritance for both markers and the standard deviation are calculated as well. The data is subdivided in the following file tabs:

1. **Fig. 6 (Original):** Inheritance of the *w5*-CopyCat targeting the *white* gene carrying the “Original” gRNA scaffold variant, driven by a transgene expressing Cas9 under the *vasa* promoter.
2. **Fig. 6 (Loop):** Inheritance of the *w5*-CopyCat carrying the “Loop” gRNA scaffold variant.
3. **Fig. 6 (Mutation):** Inheritance of the *w5*-CopyCat carrying the T>C “Mutation” gRNA scaffold variant.
4. **Fig. 6 (Loop+Mutation):** Inheritance of the *w5*-CopyCat carrying the gRNA scaffold variant with both the additional “Loop” as well as the T>C “Mutation”.
5. **Summary:** This tab summarizes the total of flies counted in the four experimental conditions, and reports the average values for inheritance, cutting and conversion rates observed.
6. **St: Inheritance rates statistics:** Statistical analysis comparing different inheritance rates observed. The “Loop”, “Mutation” and the “Loop+Mutation” conditions were compared to the “Original” condition.
7. **St: Cutting rates statistics:** Statistical analysis comparing different cutting rates comparison. The “Loop”, “Mutation” and the “Loop+Mutation” conditions were compared to the “Original” condition.
8. **St: Conversion efficiency statistics:** Statistical analysis comparing different conversion efficiencies comparison. The “Loop”, “Mutation” and the “Loop+Mutation” conditions were compared to the “Original” condition.

**SUPPLEMENTARY METHODS**

**Plasmid Constructs**: Plasmid sequences will be deposited to GenBank ahead of publication

**Cas9 plasmids**

1) **pVMG177 (*Cq-Actin5C*>Cas9_wHAs)**

   The 5' regulatory sequence (~3.4kb upstream) and 3' terminator sequence (~0.4kb downstream) of *Cx. quin-Act5C* gene (Ref # CPIJ009808) were amplified from the genomic DNA of *Cx. quin*. CA strain and ligated with Cas9 protein (amplified from *Aste.* Cas9 plasmid) to build *Act5C*>Cas9 plasmid (**pVMG173**). The *Act5C*>Cas9 element then ligated with *Hr5IE1*>GFP fluorescent marker with Gibson Assembly. This component was later digested with BbsI enzyme and liagted with Cquin w4 HAs backbone (~1.0kb homology arms around the white-sgRNA-4 cutting site). The white gene (Ref # CPIJ005542) is amplified from *C. quin*. CA strain.

   Primers used in this construct: 494, 495, V0915, V0889, V0888, V0914, V0700, V0638, V0882, V0883, V0325, V0884, V0885, 941, V0887, V0886.

2) **pVMG220** (***Cq-Rpl40*>Cas9_wHAs**)

   The 5' regulatory sequence (~3.4kb upstream) and 3' terminator sequence (~0.9kb downstream) of *Rpl40* gene (Ref # CPIJ002413) were amplified from the genomic DNA of *C. quin*. CA strain and ligated with Cas9 protein (amplified from *Aste.* Cas9 plasmid) to build Rpl40>Cas9 plasmid (**pVMG193**). The Act5C>Cas9 element then ligated with *Hr5IE1*>GFP fluorescent marker with Gibson Assembly. This component was later ligated with *C. quin* w4 HAs backbone.

   Primers used in this construct: V0834, V0832, V0963, V0964, V0325, V0884, V1044, V1040, V0512, V0408, V1068, V1069, V1000, V1070.

3) **pVMG235 (*Cq-nanos*>Cas9_wHAs)**

   The 5' regulatory sequence (~3.3kb upstream) and 3' terminator sequence (~1.4kb downstream) of *nanos* gene (Ref # CPIJ011551) were amplified from the genomic DNA of *C., quin*. CA strain and ligated with Cas9 protein (amplified from *Aste.* Cas9 plasmid) to build *nanos*>Cas9 plasmid (**pVMG212**). The *nanos*>Cas9 element then ligated with Opie2>DsRed fluorescent marker with Gibson Assembly. This component was later ligated with *C. quin* w4 HAs backbone.

   Primers used in this construct: V0660, V0678, V1009, V1011, V0325, V0884, V0982, V0656, V1010, V1119, V0754, V1118, V1117, V1068, V1062, V0335, V1029.

4) **pVMG236 (*Cq-vasa*>Cas9_wHAs)**

   The 5' regulatory sequence (~2.4kb upstream) and 3' terminator sequence (~0.8kb downstream) of *vasa* gene (Ref # CPIJ009286) were amplified from the enomic DNA of *C. quin*. CA strain and ligated with Cas9 protein (amplified from *Aste.* Cas9 plasmid) to build *vasa*>Cas9 plasmid (**pVMG213**). The *vasa*>Cas9 element then ligated with Opie2>DsRed fluorescent marker with Gibson Assembly. This component was later ligated with *C. quin* w4 HAs backbone.

   Primers used in this construct: V0957, V0981, V0325, V0884, V1013, V0540, V1014, V1012, V0957, V1121, V0754, V0335, V1029, V1062, V1028, V0754, V1120, V1117, V1068.

5) **pVMG225 (*IE1*>Cas9-P2A-DsRed)**

The 5' regulatory sequence (~1.1kb upstream) and 3' terminator sequence (~0.2kb downstream) of *Hr5IE1* gene were amplified from Baculovirus and ligated with Cas9 protein and T2A>DsRed using Gibson Assembling.

Primers used in this construct: V0887, V0886, V1060, V1059, 1167, V0065, V0053, V1061.

**U6>gRNA plasmids**

1) **pVMG146 (*Cq*-U6:1>2XBbsI-gRNA)**

The promoter and terminator sequences of U6:1 (CPIJ039653) were amplified from *Cquin*. AL line, and ligated with a double-BbsI restriction site linker for later insertion of different gRNAs.

Primers used in this construct: V0621, V0622, V0623, V0624.

2) **pVMG147 (*Cq*-U6:2a>2XBbsI-gRNA)**

The promoter and terminator sequences of U6:2a (CPIJ039728) were amplified from *Cquin*. AL line, and ligated with a double-BbsI restriction site linker for later insertion of different gRNAs.

Primers used in this construct: V0522, V0628, V1412, V0693.

3) **pVMG217 (*Cq*-U6:2b>2XBbsI-gRNA)**

The promoter and terminator sequences of U6:2b (CPIJ039728) were amplified from *Cquin*. CA line, and ligated with a double-BbsI restriction site linker for later insertion of different gRNAs.

Primers used in this construct: V0575, V0462, V1032, V1034.

4) **pVMG218 (*Cq*-U6:4>2XBsmBI-gRNA)**

The promoter and terminator sequences of U6:4 (CPIJ039801) were amplified from *Cquin*. CA line, and ligated with a double-BsmBI restriction site linker for later insertion of different gRNAs.

Primers used in this construct: V0465, V0466, V1035, V1036.

5) **pVMG149(*Cq*-U6:6>2XBbsI-gRNA)**

The promoter and terminator sequences of U6:6 (CPIJ039596) were amplified from *Cquin*. AL line, and ligated with a double-BbsI restriction site linker for later insertion of different gRNAs.

Primers used in this construct: V0695, V0696, V0697, V0698.

6) **pVMG164 (*Cq*-U6:7>2XBbsI-gRNA)**

The promoter and terminator sequences of U6:7 (CPIJ040693) were amplified from *Cquin*. CA line, and ligated with a double-BbsI restriction site linker for later insertion of different gRNAs. This plasmid was synthesized by IDT-Geneblock based on the amplified sequence.

Primers used in this construct: V0533, V0534.

The *cd1*-sgRNA and *Kh3*-sgRNA were synthesized and later replaced the BbsI restriction site to build different U6-gRNA plasmids for experiments.

Anneal oligos for *cd1*: V1209 and V1212

Anneal oligos for *kh3*: V1141 and V1142

**HDR templates for *Culex* transgenesis**

**1) pVMG252 (*Cq-vasa*>Cas9_cdHAs_O)**

The *vasa*>Cas9 element was amplified from **pVMG213** then ligated with Opie2>DsRed marker, and this transgene later inserted between two ~1.5 kb homology arms (HAs) matching the genomic sequences of the *cardinal* locus (CPIJ005949) abutting the *cd1*-gRNA target site. The U6:1>cd1-gRNA component was placed outside of the HAs.

Primers used in this construct: V1181, V1182, V1185, V1184, V0754, V1062, V1186, V1120, V0540, V0957, 941, V1219, V0624, V1227.

**2) pVMG280 (*Cq-vasa*>Cas9_cdHAs_L)**

The HAs and transgene are the same as **pVMG252**, the difference is the gRNA scaffold with a modified "Loop" structure.

Primers used in this construct: V1181, V1182, V1185, V1184, V0754, V1062, V1186, V1120, V0540, V0957, V1263, V1264, V0624, V1227, 941, V1219.

**3) pVMG277 (*Cq-vasa*>Cas9_cdHAs_L+M)**

The HAs and transgene are the same as **pVMG252**, the difference is the gRNA scaffold with a modified "Loop+Mutation" structure.

Primers used in this construct: V1181, V1182, V1185, V1184, V0754, V1062, V1186, V1120, V0540, V0957, V1263, V1267, V0624, V1227, 941, V1219.

**Copy-Cat constructs used in *Drosophila* experiments**

**1) pVMG109 (CC-U6:3-*w5*-GFP-wHAs_O)**

The *Dmel*-U6:3>w5 were synthesized by GeneBlock (IDT), and ligated with the 3XP3 driving EGFP fluorescent marker. These components were later inserted into the *drosophila white* gene at w5 cutting site with ~1.0kb homology arms.

Primers used in this construct: 451, 611, V0308, V0309, V0358, V0359, 1068, V0357.

**2) pVMG221** (**CC-U6:3-*w5*-GFP-wHAs_L**)

This construct was built based on the **pVMG109** with a "Loop" modification on its scaffold structure.

Primers used in this construct: V0359, V1049, V1048, V0897, V1050, 1068.

**3) pVMG222** (**CC-U6:3-*w5*-GFP-wHAs_M**)

This construct was built based on the **pVMG109** with a T>C "Mutation" modification on its scaffold structure.

Primers used in this construct: V0359, V1049, V1048, V0897, V1050, 1068, V1051, V1116.

**4) pVMG274** (**CC-U6:3-*w5*-GFP-wHAs_L+M**)

This construct was built based on the **pVMG221** with both a C>T “Mutation” and “Loop” modifications on its scaffold structure.

Primers used in this construct: V1051, V1116, V1267, V1268.

#### Primer List

| Name | Primer |
|---|---|
| **Primers used for building constructs** | |
| 494 | GGTATCAGCTCACTCAAAGGCGGTAATACGG |
| 495 | GAAAGGGCCTCGTGATACGCCTATTTTTATAGG |
| V0915 | AATAGGCGTATCACGAGGCCCTTTCGACTAACGTACCAATGCACCTGG |
| V0889 | GAAGACAGGCGAAAGCCAAACGTTTACGCTCACCTCCCGACAGACCCTTGATCCGTCCTG |
| V0888 | CGTTTGGCTTTCGCctGTCTTCatctaGAAGACatCCGAGAGTGCGTCGTTCATCAAATAGGCGATCGCGGTGG GGAGAGGAAACGTCTG |
| V0914 | GCCTTTGAGTGAGCTGATACCATTGCAAGTTCCTAACCATAACCTAC |
| V0700 | GGCCAATTTGCCAATTTACCCCCG |
| V0638 | CTATTTGATGAACGACGCACTCTCGG |
| V0882 | CGAAGAAGAAGCGCAAGGTGTAATAAGCGCCTTCACGCCGCAACAAC |
| V0883 | TCCAGACCGATCGAGTACTTCTTATCCATGTTGATATCTGCGCGCAAG |
| V0325 | GATAAGAAGTACTCGATCGGTCTGGATATC |
| V0884 | TTACACCTTGCGCTTCTTCTTCG |
| V0885 | CGAGAGTGCGTCGTTCATCAAATAGGCGATCGCgGAGACCGCTCACTCAAAGGCGGTAATACGG |
| 941 | GGGCGAATTCTGCAGATATCCATCAC |
| V0887 | GGTAAATTGGCAAATTGGCCGCGATCGCttaGGCGCGCCcgcgtaaaacacaatcaagtatg |
| V0886 | GATGGATATCTGCAGAATTCGCCCTTGGTCTCaGGCGCGCCGGTCGACTGATCATAATCAGC |
| V0834 | GCATTTTCTGCCTGCTTCTTTTCTCACGG |
| V0832 | GTCATCGTGGTAAGCGAATCTAG |
| V0963 | CCGAAGAAGAAGCGCAAGGTGTAAGcCCCCATTCGTTCGCCGTTTG |
| V0964 | CCGATCGAGTACTTCTTATCCATGGTGAATCTGCGGGAACGGAAAG |
| V1044 | GTTTCCTCTCCCCACCGCGATCGCGTCATCGTGGTAAGCGAATCTAG |
| V1040 | CGCGGGCGCGCCtaaGCGATCGCGCATTTTCTGCCTGCTTCTTTTC |
| V0512 | GCATACAGAGCTTAGCTCTGTTC |
| V0408 | GATCGAATGCAATAATCACCACGTAAG |
| V1068 | GATCGCGGTGGGGAGAGGAAACGTCTGG |
| V1069 | GAAAGCCAAACGTTTACGCTCACCTCCCGACAGACCCTTGATCCGTCCtG |
| V1000 | GTGAGCGTAAACGTTTGGCTTTC |
| V1070 | atcgcttaGGCGCGCCcgcg |
| V0660 | ATGTCCTTTTTAATCACGATTCAC |
| V0678 | GGCTTTTTTACGAAAACCTCCAACTC |
| V1009 | GACCGATCGAGTACTTCTTATCCATGTTTTAGAAAGGACCTTGcCGGG |
| V1011 | ACAAATTTTGAGCAACCTTGCTCAAGGGCGAATTCTGCAGATATCC |
| V0982 | AATTGTAGTAATTTGAACCATGTAAGAG |
| V0656 | GAGCAAGGTTGCTCAAAATTTGTCTG |
| V1010 | GAAGAAGAAGCGCAAGGTGTAAAATTGTAGTAATTTGAACCATGTAAGAG |
| V1119 | CCAGACGTTTCCTCTCCCCACCGCGATCGAGCAAGGTTGCTCAAAATTTGTCTG |
| V0754 | TTAAGATACATTGATGAGTTTGGACAAACCACAACTAGAATGC |
| V1118 | GTGAATCGTGATTAAAAAGGACATtcggcatactcggtggcctcccc |

| | |
|---|---|
| V1117 | GTCCAAACTCATCAATGTATCTTAacgttaactcgaatcgctatccaagc |
| V1062 | TCGGcatactcggtggcctcc |
| V0335 | TAAGATACATTGATGAGTTTGGACAAACCACAACTAGAATGCAGTGAAAAAAATG |
| V1029 | ccgcaacctgtctctggtgatgGCCTCCTCCGAGGACG |
| V0981 | GTTGGCGTTGACTTACTGAAAAG |
| V0957 | CCGTGCATAATTCGATTTGATGCG |
| V1121 | CCAGACGTTTCCTCTCCCCACCGCGATCGCGCTCTTCaGCTAGCCCACCCAATGTAC |
| V0754 | TTAAGATACATTGATGAGTTTGGACAAACCACAACTAGAATGC |
| V1120 | CGCATCAAATCGAATTATGCACGGtcggcatactcggtggcctccccac |
| V1060 | CGCCACCACCTGTTCCTGTAGACATTGATGAGTTTGGACAAACCAC |
| V1059 | CCGTCGTGGTCCTTATAGTCCATgtcacttggttgttcacgatcttg |
| V1167 | ATGGACTATAAGGACCACGACGG |
| V0065 | CCGAACAGGCCATTCTTCTTCTC |
| V0053 | GAGAAGAAGAATGGCCTGTTCGG |
| V1061 | CTACAGGAACAGGTGGTGGCG |
| V0621 | GCAAGTTAAAATAAGGCTAGTCCGTTATCAACTTGAAAAAGTGGCACCGAGTCGGTGCATTTTTTTGGTGGC GGATGAGTAACAACATTC |
| V0622 | CTTGCAAGTCGTCGTCGTCGTCTGGGAAGGGCGAATTCTGCAGATATC |
| V0623 | CGGACTAGCCTTATTTTAACTTGCTATTTCTAGCTCTAAAACAGGTCTTCTCGAAGACCCAAGTTCGGAGGGT GCCCTCGTTGCTTATAT |
| V0624 | CCAGACGACGACGACGACTTGCAAG |
| V0522 | CAGTGCTGCCAGATTTTCGATAAAC |
| V0628 | GGAGGCTCATTTCCTTAAACTCGTG |
| V1412 | TCTAAAACGGGTCTTCGAGAAGACCTGAGTACAAAGGGCTCAAAATTTG |
| V0693 | GCTAGAAATAGCAAGTTAAAATAAGGCTAGTCCGTTATCAACTTGAAAAAGTGGCACCGAGTCGGTGCTTTTT TTTTGATTCTGG |
| V0575 | GCCGAAGCAAAGACAAGTTC |
| V0462 | GTTTTACGGTGCACATCAATCTGAG |
| V1032 | GTCCGTTATCAACTTGAAAAAGTGGCACCGAGTCGGTGCTTtTtttttGATTCTGGTTTaTTTTTGTTcTCTTTTGG AAG |
| V1034 | TAGCCTTATTTTAACTTGCTATTTCTAGCTCTAAAACAGGTCTTCTCGAAGACCCGAGTACaaaGGGCTCAAAa TTTGCTTATATAG |
| V0465 | GTATCCGTTGGCTAGTAGAAAAACTC |
| V0466 | GCACTTCATTGCCATTCACAATAG |
| V1035 | CCTTATTTTAACTTGCTATTTCTAGCTCTAAAACAGAGACGATCGTACGTCTCGAAaTTAtGTTGTCTACATATTT GCTTATATAGTTAG |
| V1036 | CTAGTCCGTTATCAACTTGAAAAAGTGGCACCGAGTCGGTGCTTTtttttGGCCatAAACAATTTAAaTCG |
| V0695 | GTTAAAATAAGGCTAGTCCGTTATCAACTTGAAAAAGTGGCACCGAGTCGGTGCTTTTTTTTTAGGGCGAATT CTGCAGATATCCATCAC |
| V0696 | GTAGACCTTATTATCTTCGTAGATTTTGAAGGGCGAATTCCAGCAC |
| V0697 | CAAAATCTACGAAGATAATAAGGTCTACTATTAC |
| V0698 | TTGATAACGGACTAGCCTTATTTTAACTTGCTATTTCTAGCTCTAAAACAGGTCTTCTCGAAGACCCAAGTTGT AGGGCCGGGAAAGTGC |
| V0533 | GCTCAAATTTTTGTGGGAGTTCCTTG |
| V0534 | GGTTTTGGTAGAGAGTTGCTCTCTTATG |
| V1181 | CTTGCTGTTTGGGGTCAATTCTTG |
| V1182 | GTTATATATGGCCCATCAGGCTCG |
| V1184 | ggaggccaccgagtatgCCGATTGCGGCAGATGATGCTGGAC |
| V1185 | GTCCAAACTCATCAATGTATCTTAACTCGGATGCCATCAGCCATTCCCCG |
| V1186 | GTACATTGGGTGGGCTAGCtGAAGAGCTTGCGGCAGATGATGCTGGACAGTTTC |

| | |
|---|---|
| V0540 | GCTCTTCaGCTAGCCCACCCAATGTACCATAGTGCAAC |
| V0957 | CCGTGCATAATTCGATTTGATGCG |
| 941 | GGGCGAATTCTGCAGATATCCATCAC |
| V1219 | CTTGCAAGTCGTCGTCGTCGTCTGGGTTATATATGGCCCATCAGGCTCG |
| V1263 | CAACTCGGATGCCATCAGCCAAGTTCG |
| V1267 | GTTTcAGAGCTAtgctgGAAAcagcaTAGCAAGTTgAAATAAGGCTAGTCCGTTATCAAC |
| V1227 | GTGATGGATATCTGCAGAATTCGCCCTAAGGAACACCTGTTCCCTCTGCGAG |
| 451 | GTACGCGTATCGATAAGCTTtaaGATACATTGATGAGTTTGG |
| 611 | CTGCGGCGATCGAAAGGCAAGGGCATTCAGC |
| V0308 | CTTTCGATCGCCGCAGacgtcgtaaggctagttttgagaaggatcgcttgtctgggcaag |
| V0309 | ATGTATCttaAAGCTTATCGATACGCGTACgctagccaaggtgctacgaaatccgttgtg |
| V0358 | ctcaaaactagccttacgacgtGGCGATACTTGGATGCCCTGCGGCGATCGAAAGGCAAG |
| V0359 | ATCCCCGGGCGAGCTCGcatatATCCGGGATGCGACTGCTCAATG |
| 1068 | atatgCGAGCTCGCCCGGGGATC |
| V1049 | AtgctgTTTCcagcaTAGCTCTAAAACGATGGCGATACTTGGATGCCgacgttaaattg |
| V0357 | acgtcgtaaggctagttttgagaaggatcgcttgtctgggcaag |
| V1048 | TAGAGCTAtgctgGAAAcagcaTAGCAAGTTAAAATAAGGCTAGTC |
| V0897 | gggccGCGACTCTAGATCATAATCAGCCATACCACAT |
| V1050 | GATTATGATCTAGAGTCGCGGCCcCTATTACTTGTACAGCTCGTCCATG |
| V0359 | ATCCCCGGGCGAGCTCGcatatATCCGGGATGCGACTGCTCAATG |
| V0897 | gggccGCGACTCTAGATCATAATCAGCCATACCACAT |
| V1051 | TCGTTTcAGAGCTAGAAATAGCAAGTTgAAATAAGGCTAGTCCGTTATCAAC |
| V1116 | TGGCGATACTTGGATGCCgacgttaaattgaaaatagg |
| V1267 | GTTTcAGAGCTAtgctgGAAAcagcaTAGCAAGTTgAAATAAGGCTAGTCCGTTATCAAC |
| V1268 | GATGGCGATACTTGGATGCCgacgttaaattg |
| **Anneal Oligos for *Kh-3* gRNA** | |
| V1141 | aaacACCGAGTTTCGCTAAACTCC |
| V1142 | ACTCGGAGTTTAGCGAAACTCGGT |
| **Anneal Oligos for *cd-1* gRNA** | |
| V1209 | AAACCAACTCGGATGCCATCAGCC |
| V1212 | ACTCGGCTGATGGCATCCGAGTTG |
| **Primers used for *cd-1* site Deep sequencing (amplicon added, marker as green)** | |
| V1217 | ACACTCTTTCCCTACACGACGCTCTTCCGATCTCCGCTGATCACGTGCCTTCTCGG |
| V1218 | GACTGGAGTTCAGACGTGTGCTCTTCCGATCTGCACTTGTTCCCACGCTGGCTTC |
| **Primers used for *Kh-3* site Deep sequencing (amplicon added, marker as green)** | |
| V1157 | ACACTCTTTCCCTACACGACGCTCTTCCGATCTTGGTCATGATCAAGTGCAAACCG |
| V1158 | GACTGGAGTTCAGACGTGTGCTCTTCCGATCTAAGGAAACGGAGTTGTACAGCG |
| **Primer used for NGS analysis in Hsu cells (*Kh-3* target site)** | |
| NGS218 | ATGCGGCACACGCCATGGTT |
| NGS219 | ACGAGTCTTCCTACCTCGATGTAGTTGTA |
| **Primers used to isogenize at the *cardinal* locus** | |
| V0917 | GACCCTGATGAAACAAACACGC |
| V0918 | CTTCAAGTGGCTGGCACCG |
